# Structure-based probe reveals the presence of large transthyretin aggregates in plasma of ATTR amyloidosis patients

**DOI:** 10.1101/2024.03.09.584228

**Authors:** Rose Pedretti, Lanie Wang, Anna Yakubovska, Qiongfang S. Zhang, Binh Nguyen, Justin L. Grodin, Ahmad Masri, Lorena Saelices

## Abstract

ATTR amyloidosis is a relentlessly progressive disease caused by the misfolding and systemic accumulation of amyloidogenic transthyretin into amyloid fibrils. These fibrils cause diverse clinical phenotypes, mainly cardiomyopathy and/or polyneuropathy. Little is known about the aggregation of transthyretin during disease development and whether this has implications for diagnosis and treatment. Using the cryogenic electron microscopy structures of mature ATTR fibrils, we developed a peptide probe for fibril detection. With this probe, we have identified previously unknown aggregated transthyretin species in plasma of patients with ATTR amyloidosis. These species are large, non-native, and distinct from monomeric and tetrameric transthyretin. Observations from our study open many questions about the biology of ATTR amyloidosis and reveals a potential diagnostic and therapeutic target.

## Introduction

ATTR amyloidosis is a fatal disease resulting from the systemic deposition of amyloid fibrils composed of transthyretin (TTR when functional, or ATTR when in its amyloidogenic form), ultimately leading to progressive organ failure and death^1^. ATTR deposition starts with the destabilization of the functional tetrameric form of transthyretin that associates with aging in wild-type ATTR (ATTRwt) amyloidosis or a mutation in the transthyretin gene in variant ATTR (ATTRv) amyloidosis^1^. ATTRwt amyloidosis usually presents with restrictive cardiomyopathy, often with orthopedic manifestations such as bilateral carpal tunnel syndrome and/or spinal stenosis^1^. ATTRv amyloidosis has a clinical presentation that is often variable and may involve polyneuropathy, carpal tunnel syndrome, gastrointestinal and eye involvement, with or without cardiomyopathy^1^. The molecular mechanisms behind this phenotypic variability are yet to be revealed.

While it is understood that transthyretin tetramer dissociation is the rate limiting step of ATTR pathogenesis^2^, little is known about the nature of the first amyloid species that lead to disease. A recent study describes the presence of non-native transthyretin (NNTTR) species in plasma of neuropathic ATTR amyloidosis patients, which are proposed to be intermediate species during the aggregation process. These species were identified with a novel detection probe that intercalates into oligomeric species of ATTR^3^. In a follow-up study, the authors designed a novel immunoassay to recognize misfolded oligomeric ATTR species in plasma of neuropathic ATTRv amyloidosis patients^4^. These NNTTR species were not found in individuals with cardiomyopathy-associated genotypes and their presence was limited in individuals that presented with both cardiomyopathy and neuropathy. It is unclear whether cardiomyopathic ATTR amyloidosis patients have other types of circulating aggregates or intermediates in their plasma. If they do exist, they may be structurally or otherwise distinct from those NNTTR species detected in patients with a neuropathic phenotype. These early steps of the aggregation of transthyretin must be further explored to understand the biology of ATTR amyloidosis, which may facilitate the identification of new targets for therapeutic and diagnostic development.

Our laboratory has previously determined cryogenic electron microscopy (cryo-EM) structures of ATTR fibrils extracted from cardiac tissue of ATTRwt and ATTRv amyloidosis patients. These structures have revealed a common core with local conformational changes^5^.

One commonality shared among all fibril structures is the presence of β-strands F and H on the exterior of the fibril surface^5^. In previous studies, we showed that strands F and H are critical for transthyretin aggregation *in vitro*^6-9^. We hypothesize that these strands have a relevant role during transthyretin misfolding and/or aggregation in the early steps of protein aggregation.

In the present study, we use the structures of β-strands F and H in crystal and cryo-EM structures of ATTR fibrils to develop a first-generation *<u>T</u>ransthyretin <u>A</u>ggregation <u>D</u>etection* (TAD1) probe for detection of ATTR aggregates and fibrils. Our structure-specific probe binds recombinant aggregates, ATTR fibrils present in tissue, and large aggregates present in plasma of cardiac ATTR amyloidosis patients, which we have recently discovered and we further characterize here^10^. Our probe, however, does not bind to tetrameric or monomeric transthyretin. Our findings are significant as they inform about the biopathology of ATTR aggregation and provide insight into the development of ATTR amyloidosis.

## Materials and Methods

### Peptide design and synthesis

Peptide development was performed by rational design starting from our previously published peptide inhibitors that target the two amyloid-driving segments of transthyretin^7^. All sequences are listed in Supplemental Tables 1 and 2. Fluorescent and epitope modifications were added to peptides for detection purposes. Peptides were synthesized by LifeTein LLC in lyophilized form. Before running experiments, the peptide was reconstituted in distilled water to yield a final concentration of 5 mM. The resuspended peptide was filtered through a 0.22 µm filter tube by spinning at 20,000 × g for 5 minutes at 4 °C to remove any possible aggregates from the solution.

### Patients and tissue material

This study required three types of human-derived specimens: fresh-frozen cardiac tissue, serum, and plasma. Details about cardiac tissues from ATTR amyloidosis patients carrying wild-type ATTR (n=1) or variant ATTR (n=2) are listed in Supplemental Table 3. Details about serum and plasma samples of ATTR amyloidosis patients (n=20) and controls (n=11) are listed in Supplemental Table 4. Cardiac tissues from the left ventricle of either explanted or autopsied hearts were obtained from the laboratory of the late Dr. Merrill D. Benson at the University of Indiana. Serum and plasma specimens were obtained from Dr. Ahmad Masri at Oregon Health and Science University and the Dallas Heart Study at UTSW^11^. The Office of the Human Research Protection Program at UTSW granted exemption from Internal Review Board review because all specimens were anonymized. All experiments involving blood samples were blinded to the researchers running and analyzing the assays.

### Extraction of amyloid fibrils from human cardiac tissue

*Ex vivo* fibrils were extracted from fresh-frozen heart tissue as described by Nguyen, Afrin, Singh, et al^5^. Briefly, ∼100 mg of frozen cardiac tissue per patient was thawed at room temperature and minced into small pieces with a scalpel. The minced tissue was suspended in 400 µL Tris-calcium buffer (20 mM Tris, 150 mM NaCl, 2 mM CaCl_2_, 0.1% NaN_3_, pH 8.0) and centrifuged for 5 minutes at 3,100 × g and 4 °C. The pellet was washed and centrifuged in Tris-calcium buffer three additional times. After washing, the pellet was resuspended in 375 µL of 5 mg/mL collagenase (Sigma Aldrich C5138) in Tris-calcium buffer. This solution was incubated overnight at 37 °C with shaking at 400 rpm. The resuspension was centrifuged for 30 minutes at 3100 × g and 4 °C and the pellet was resuspended in 400 µL Tris-ethylenediaminetetraacetic acid (EDTA) buffer (20 mM Tris, 140 mM NaCl, 10 mM EDTA, 0.1% NaN3, pH 8.0). The suspension was centrifuged for 5 minutes at 3,100 × g and 4 °C, and the washing step with Tris–EDTA was repeated nine additional times. Following washing, the pellet was resuspended in 200 μL ice-cold water supplemented with 5 mM EDTA and centrifuged for 5 minutes at 3,100 × g and 4 °C. This step released the amyloid fibrils from the pellet, which were collected in the supernatant. EDTA helped solubilize the fibrils. This extraction step was repeated five additional times. The material from the various patients was handled and analyzed separately.

### Preparation of fibril seeds

Extracted fibrils were treated with 1% sodium dodecyl sulfate (SDS) in sodium phosphate-EDTA buffer, and the soluble fraction was discarded after centrifugation at 13,000 rpm for 20 minutes. This process was repeated twice. The sample was then washed with sodium phosphate-EDTA buffer (without SDS) three times by centrifugation. Following the washes, the sample was sonicated in a bath sonicator, with cycles of 5 seconds on/5 seconds off, for a total of 10 minutes using an amplitude of 85 (Q700 sonicator, Qsonica). The protein concentration was measured using the Pierce Micro BCA Protein Assay Kit (Thermo Fisher Scientific 23235).

### Thioflavin T inhibition assay

We used a thioflavin T (ThT) fluorescence assay to characterize the inhibition profile of peptides following established protocols^6,8^. Briefly, each 60 µL reaction contains 0.5 mg/mL recombinant monomeric transthyretin (MTTR), 10% phosphate buffered saline, 5 µM ThT, and 30 ng/mL seeds, along with the stated peptide concentration. Unless otherwise stated, each reaction was triplicated and pipetted into 384 well plates (Thermo Scientific 242764). The plates were then incubated at 37 °C for 132 hours shaking at 700 rpm. ThT fluorescence emission was measured at 482 nm with excitation at 440 nm in a FLUOstar Omega (BMG LabTech) microplate reader.

### Negative-stained transmission electron microscopy

Samples of interest after thioflavin screening were observed as previously described^7^. Briefly, a 5 µL sample was spotted onto a glow-discharged carbon film 300 mesh copper grid (Electron Microscopy Sciences CF300-Cu-50), incubated for two minutes then gently blotted onto filter paper to remove excess solution. The grid was negatively stained with 5 µL of 2% uranyl acetate for two minutes and gently blotted to remove the solution. Another 5 µL of 2% uranyl acetate was applied on the grid and immediately blotted away. Samples were examined using an FEI Tecnai 12 electron microscope at an accelerating voltage of 120 kV.

#### In vitro *aggregation assay of transthyretin*

Generation of recombinant transthyretin aggregates was done as described previously^7^. Briefly, 1 mg/mL of tetrameric transthyretin with an N-terminal polyhistidine tag was incubated in aggregation buffer containing 10 mM sodium acetate (pH 4.3), 100 mM KCl, and 1 mM EDTA at 37 °C for four days. At the specified time points, 35 µL of the reaction was removed, and the total transthyretin content and ATTR were estimated using anti-transthyretin and TAD1 immunodotblotting, respectively. For anti-transthyretin immunodotblotting, 5 µL of sample was dotted onto 0.2 µm nitrocellulose membrane (BioRad 1620112) and blocked for 30 minutes with 10% milk in 10% tris-buffered saline, 0.1% Tween-20 (TBS-T). After washing the membrane three times for five minutes with 1X TBS-T, the membrane was incubated with an anti-transthyretin antibody (1:1,000; Genscript) overnight in 5% milk in TBS-T. The next day, the membrane was washed three times for ten minutes and further incubated with goat anti-rabbit secondary antibody (1:1,000; Invitrogen 31460) for one hour. The membrane was washed three times for ten minutes and then incubated for 5 minutes with an enhanced chemiluminescence reagent (Promega W1001). The blot was imaged in an Azure Biosystems C600 imaging system. The protocol for TAD1 immunodotblotting was the same but with the following modifications: the membrane was blocked in TBS-T buffer containing 10% bovine serum albumin (BSA); and incubated with 5 µM TAD1 in 10% BSA/TBS-T. After washing unbound peptide, the fluorescence intensity of TAD1 binding was measured in an Azure Biosystems C600 imaging system through excitation of the membrane at 472 nm and reading emission at 513 nm.

Additionally, 50 µL of the reaction was collected at each specified time and spun at 13,000 rpm for 30 minutes to pellet insoluble aggregates. The pellet was then resuspended in 50 µL fresh aggregation buffer, and centrifuged for the second time. Eventually, the pellet was resuspended in 50 µL 6 M guanidine hydrochloride. 2 µL of this mixture was diluted 10 times with guanidine hydrochloride before analysis. The polyhistidine tag in the insoluble fraction was probed with the SuperSignal West HisProbe Kit (ThermoFisher Scientific 15168) according to the manufacturer’s recommendation with a modification: the membrane was probed with 1:20,000 working solution.

#### *Immunogold labeling of* ex vivo *fibrils*

5 µM TAD1 was mixed with 50 nM Ni-NTA Nanogold (Nanoprobes 2082) in binding buffer (20 mM Tris pH 7.6, 150 mM NaCl) overnight at 4 °C. The next day the nanogold beads pelleted to the bottom of the reaction mixture. The supernatant containing unbound nanogold beads was removed. 1 µg/mL of fibrils extracted from an ATTRwt amyloidosis patient was spiked into the pellet and the solution was loaded onto a glow-discharged carbon-coated electron microscopy grid. The grid was prepared and visualized as described above.

#### Native gel electrophoresis of extracted fibrils

Native gel electrophoresis was conducted using the NativePAGE Novex Bis-Tris System (Invitrogen BN1002BOX). 5 µg of *ex-vivo* ATTR cardiac fibrils or 5 µg of recombinant protein was mixed with NativePAGE 4x sample buffer according to the manufacturer’s recommendation to a final volume of 12 µL. 10 µL of each sample was loaded into a well of NativePAGE 4-12% Bis-Tris Gels (Invitrogen NP0321BOX). The gels were run at 150 V for approximately two hours at 4 °C and transferred onto 0.2 µm PVDF membranes (Millipore IPVH00010) using the Mini Trans-Blot cell system (BioRad 1703930) for one hour at 25 V. Proteins were fixed to the membrane by incubation with 8% acetic acid for 15 minutes. The membranes were then subjected to the same staining protocol as described earlier, with the following modifications: the membranes were incubated overnight with either a primary anti-transthyretin antibody (1:1,000; Genscript) or 5 µM TAD1.

#### Denaturing Western blot of extracted fibrils

1 µg of recombinant protein or 5 µg of *ex-vivo* ATTR cardiac fibrils were boiled at 95 °C for 10 minutes. The samples were loaded onto three independent SurePAGE Bis-Tris 10x8 4-12% gels (Genscript M00652) and run at 150 V for approximately one hour. One gel was treated for immunostaining using an anti-transthyretin antibody, and the other gel was treated for the detection of fluorescence upon binding to TAD1. Gels were transferred onto nitrocellulose membranes at 25 V for 16 minutes using the Trans-Blot Turbo System (BioRad 1704150). The membranes were placed into 10% milk or 10% BSA in TBS-T for one hour to prevent non-specific binding and washed three consecutive times in TBS-T for 5 minutes. One membrane was incubated with the anti-transthyretin antibody (1:1,000; Genscript) in 5 % milk overnight, and the other membrane was incubated with 5 µM TAD1 in 10% BSA/TBS-T for one hour. The membranes were then washed three times for 10 minutes. The fluorescence intensity of TAD1 binding was measured as stated above. The membrane probed with an anti-transthyretin antibody was further incubated with goat anti-rabbit secondary antibody (1:1,000; Invitrogen 31460). The membrane was washed three times for ten minutes and then incubated with an enhanced chemiluminescence reagent (Promega W1001). The blot was imaged in an Azure Biosystems C600 imaging system.

#### Filtration assay with patient plasma

60 µL of plasma samples from ATTR amyloidosis patients and controls were subjected to filtration using a 0.22 µM centrifugal filter tube (Corning 8161). The samples were spun at 10 second intervals at 1000 g, 4 °C until there was 20 µL of filtrate at the bottom of the tube. 20 µL of unfiltered plasma, plasma that did not pass through the filter (void) and plasma that did pass through the 0.22 µM filter (filtrate) were dotted onto nitrocellulose membrane and probed with 10 µM TAD1 and anti-transthyretin antibody as described above.

#### Native gel shift assay with TAD1

30 µg of ATTRwt amyloidosis patient plasma or control plasma was incubated overnight with increasing concentrations of TAD1 (0, 12.5, 25, 50, 100 µM TAD1). The samples were then subjected to gel electrophoresis under non-denaturing conditions followed by western blotting with an anti-transthyretin antibody as described above. For quantification, the banding patterns were segmented into three groups (high molecular weight aggregates, oligomers, and tetramers) based on their associated molecular weight. These bands were quantified using ImageJ software^12^.

#### Fluorescent immunodot blotting of extracted fibrils and blood samples

The binding of TAD1 to ATTR species in patient samples was evaluated using immunodot blot analysis as described above. For assays using *ex vivo* fibrils, 0.5 µg of fibrils was dotted onto membrane. For assays using blood samples, 30 µL of sample was dotted onto the membrane. The membrane was probed with 5 µM TAD1. Fluorescence intensity was quantified using the ImageJ software. The fluorescent signal was normalized with respect to the intensity of 0.5 µg of ATTRwt fibrils.

#### Statistical analysis

Statistical analysis of TAD1 fluorescence, transthyretin aggregation, and ThT signal was conducted using the GraphPad Prism software. All samples were included in the analysis. All quantitative experiments were performed using three independent replicates and are reported as means ± standard deviation of these replicates. Statistical significance between groups was compared using a one-way ANOVA with Tukey correction, where a two-sided p-value <0.05 was considered statistically significant. Outliers in each data set were identified and removed using a Grubbs test.

## Results

### Design and validation of peptide probes

To study early species formed during transthyretin aggregation, it is essential to create specific detection methods. Based on our previous studies, we hypothesize that the strands F and H of transthyretin may play an important role during aggregation process and that they adopt an amyloidogenic structure that is present in these early species. We set out to generate peptide probes that target strands F and H in this amyloidogenic conformation. Our workflow for the design of peptide probes is shown in Figure 1A. In previous studies, we determined the amyloid structures of the β-strands F and H in isolation and in *ex-vivo* ATTR fibrils, using x-ray microcrystallography and cryo-electron microscopy, respectively^5-8^. We noted that, in all ATTR fibril structures determined to date, the β-strands F and H adopt the same conformation, similar to that shown in the crystal structure of each peptide in isolation^5-8^. Capitalizing on the strategy described in Saelices et al., 2015 JBC for the design of peptide aggregation inhibitors, we designed new *<u>T</u>ransthyretin <u>A</u>ggregation <u>B</u>inders* (TABs) that cap the tip of ATTR fibrils by interacting with both β-strands F and H (Figure 1A, Supplemental Table 1). To screen and validate our TABs, we assessed their inhibitory effect of amyloid seeding as a proxy for their capacity to bind fibrils, assuming that the binding to the tip of the fibrils would inhibit fibril polymerization (Figure 1B). Amyloid seeding is the process by which preformed amyloid fibrils can template or seed fibril polymerization, thereby accelerating fibril formation^6^. We monitored this process by using recombinant monomeric transthyretin (MTTR) and *ex-vivo* ATTRwt fibril seeds, in the presence of thioflavin T (ThT), prepared as described previously^6,8^. We found that TAB3-12 was the most potent inhibitor at a concentration of 90 µM (Figure 1B, orange) and even at sub stoichiometric ratios compared to other peptide inhibitors (Figures 1C and 1D). These data suggest that TAB3-12 was a good binder to ATTR fibril seeds, and we selected it for further modification into detection probes.

**Figure 1.**
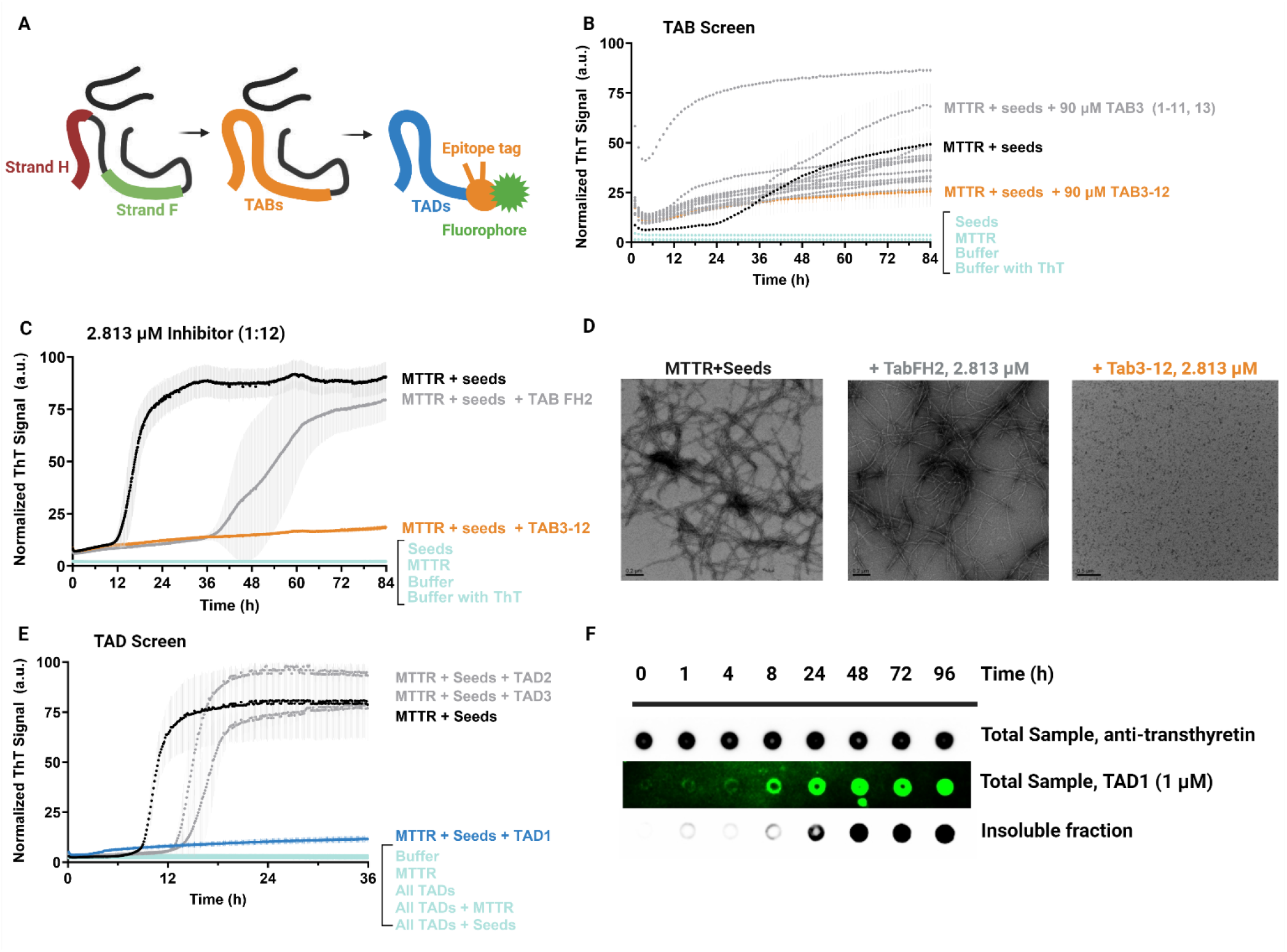
Development and screening of TAD candidates. **A**, Rational design development of peptide probes based on previously published cryo-EM structures of ATTR fibrils^5^. Fibril structure is shown top-down as black lines, with the amino acid residues corresponding to β strands F and H highlighted as the intended target for TAD peptides. **B**, ThT screening of third generation TABs to assess candidate peptide binding to fibrils. **C**, Further characterization of probe candidate TAB3-12 compared to previous generation peptide inhibitors. Peptides were added to seeding reactions in a sub stoichiometric ratio of 12 MTTR molecules to 1 peptide of interest. **D**, Electron microscopy images of ThT reactions selected from **C. E**, Assessment of TAD fibril binding activity through ThT assay. **F**, Dot blotting of recombinant transthyretin aggregation over time probing for total protein (top), aggregated species in total sample with TAD1 (middle), and insoluble components after centrifugation (bottom).

We fused TAB3-12 to three different N-terminal epitopes and a fluorescein isothiocyanate (FITC) tag, to generate first-generation *<u>T</u>ransthyretin <u>A</u>ggregation <u>D</u>etectors* (TADs) (Figure 1A, Supplemental Table 2). We confirmed that the binding of TADs to fibrils was not perturbed by the modifications using the amyloid seeding inhibition assay, as performed for TABs, and found that TAD1 (the one containing a polyhistidine tag) did not induce fibril formation, and fully inhibited seeding instead (Figure 1E). We also confirmed that the peptide did not form fibrils by itself (Figure 1E). TAD2 and TAD3 did not fully inhibit amyloid seeding (Figure 1E). We thus selected TAD1 for further studies.

To validate our structure-specific peptide design, we evaluated the binding of TAD1 to recombinant ATTR aggregates made under acidic pH using wild-type transthyretin as described previously^2,7^. With this standard procedure at pH 4.3, we dissociate the native tetrameric structure of transthyretin into unfolded monomers to promote aggregation *in vitro*. We monitored protein aggregation by collecting the insoluble fraction and visualizing the formation of aggregates by dot blotting at different time points (Figure 1F). Using an antibody that recognizes the polyhistidine tag of recombinant transthyretin, we observed that ATTR aggregates accumulated with time (Figure 1F, bottom panel), while the amount of total protein did not vary (Figure 1F, top panel). Using TAD1 fluorescence as a readout, we observed that TAD1-positive species also increase over time (Figure 1F, middle panel), confirming that TAD1 targets recombinant aggregates in a conformation-dependent manner.

### TAD1 binds ex-vivo ATTR fibrils

We visualized the localization of TAD1 binding to *ex-vivo* ATTR fibrils using nanogold beads and electron microscopy (Figure 2A). We took advantage of the polyhistidine tag present in TAD1 to coat this peptide with nickel nitrilotriacetic acid nanogold beads and confirmed that TAD1 binds to purified ATTRwt fibrils (Figure 2B). We observed indeed that the nanogold particles decorate ATTR fibrils primarily at the tips at low concentrations as well as on the surface of fibrils at higher concentrations (Figure 2B). We used tau fibrils extracted from an Alzheimer’s Disease patient as a negative control and found no TAD1 binding to tau fibrils, indicating that TAD1 is specific for fibrils made of transthyretin (Figure 2C). The sensitive binding of nanogold particles to the tip of the fibrils is consistent with our structure-based peptide design (Figure 1A).

**Figure 2.**
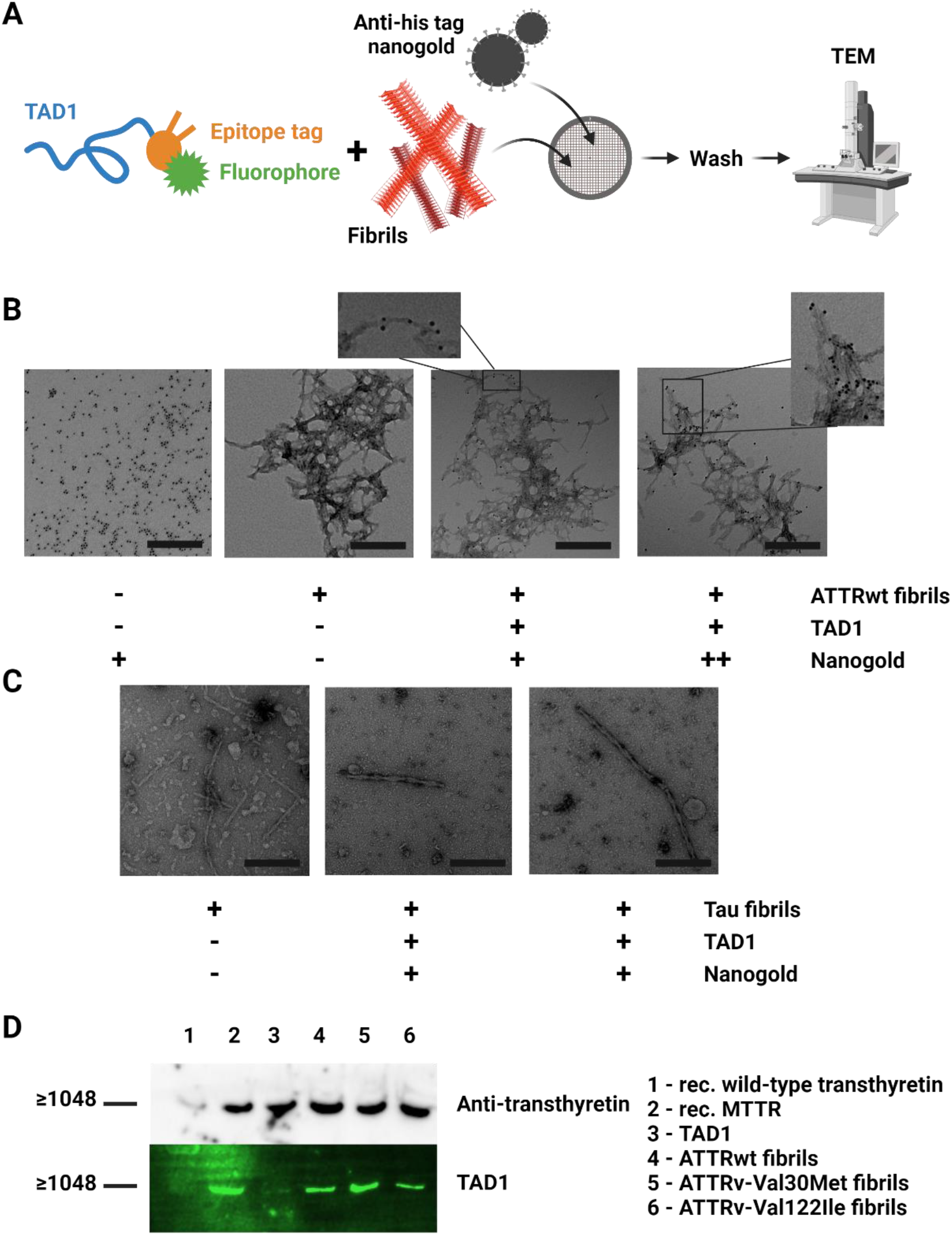
Validation of TAD1 binding to ATTR fibrils. **A**, Scheme for labeling experiments. **B**, Nanogold labeling of ATTRwt fibrils using TAD1 and either 50 µM (+) or 250 µM of nanogold (++) **C**, Nanogold labeling of tau fibrils extracted from brain using TAD1 and 50 µM nanogold. **D**, Native gel electrophoresis of TAD1, recombinant transthyretin and amyloid extracted from ATTR amyloidosis patients, blotted and probed with an anti-transthyretin antibody (top) and TAD1 (bottom).

We also observed the binding of TAD1 to *ex-vivo* fibrils by western blotting under non-denaturing and denaturing conditions (Figure 2D and Supplemental Figure 1). We observed that in non-denaturing conditions, TAD1 binds to ATTR fibrils and recombinant transthyretin aggregates that do not run through the gel. These aggregates correspond to a molecular weight higher than 1048 kDa (Figure 2D, Supplemental Figure 1A). When the same samples are subjected to denaturing conditions, TAD1 loses its ability to bind aggregates or fragments that result from denaturing ATTR fibrils (Supplemental Figure 1B), providing additional evidence that the recognition of TAD1 to ATTR species is conformational.

### TAD1 binds ATTR aggregates in plasma of cardiac ATTR amyloidosis patients

Previous studies indicate that transthyretin can adopt non-native conformations in the blood of neuropathic ATTRv amyloidosis patients, suggesting that these species could represent early stages of protein aggregation^3,4^. Since TAD1 displays high sensitivity for ATTR fibrils and not native transthyretin, we hypothesized that TAD1 could detect the presence of these non-native ATTR species (or different ones) in blood. Thus, we analyzed whether serum and plasma samples from both ATTRwt and ATTRv amyloidosis patients with cardiomyopathy or mixed phenotypes (Supplemental Table 4) contain TAD1-positive species. We first ran a pilot dot blotting assay of two patient samples and controls probing with TAD1 and found that both serum and plasma contain ATTR species that are detected by TAD1 (Figure 3A, Supplemental Figure 2). Next, we analyzed a small cohort of patient samples to investigate which sample type (serum or plasma) could provide better readouts of the presence of these species. In serum, quantifying relative fluorescence intensity of TAD1 reveals no distinct difference in binding when comparing pre-treatment patients, post-treatment patients, and controls (Figure 3B). In contrast, the analysis of plasma samples revealed a small but statistically significant difference between controls and patients pre-treatment, including both ATTRwt and ATTRv amyloidosis patients (Figure 3C). We also found a statistically significant difference in TAD1 species between the pre-treatment and post-treatment groups, regardless of the treatment type (Figure 3C). The validation of these results is now included in our recent study^10^. It is worth noting that these aggregated species are highly stable and resistant to multiple freeze thaw cycles, in contrast to the NNTTR previously described^3^ (Supplemental Figure 3). These observations suggest that there are unique ATTR species in the blood of ATTR amyloidosis patients that can be detected by TAD1.

**Figure 3.**
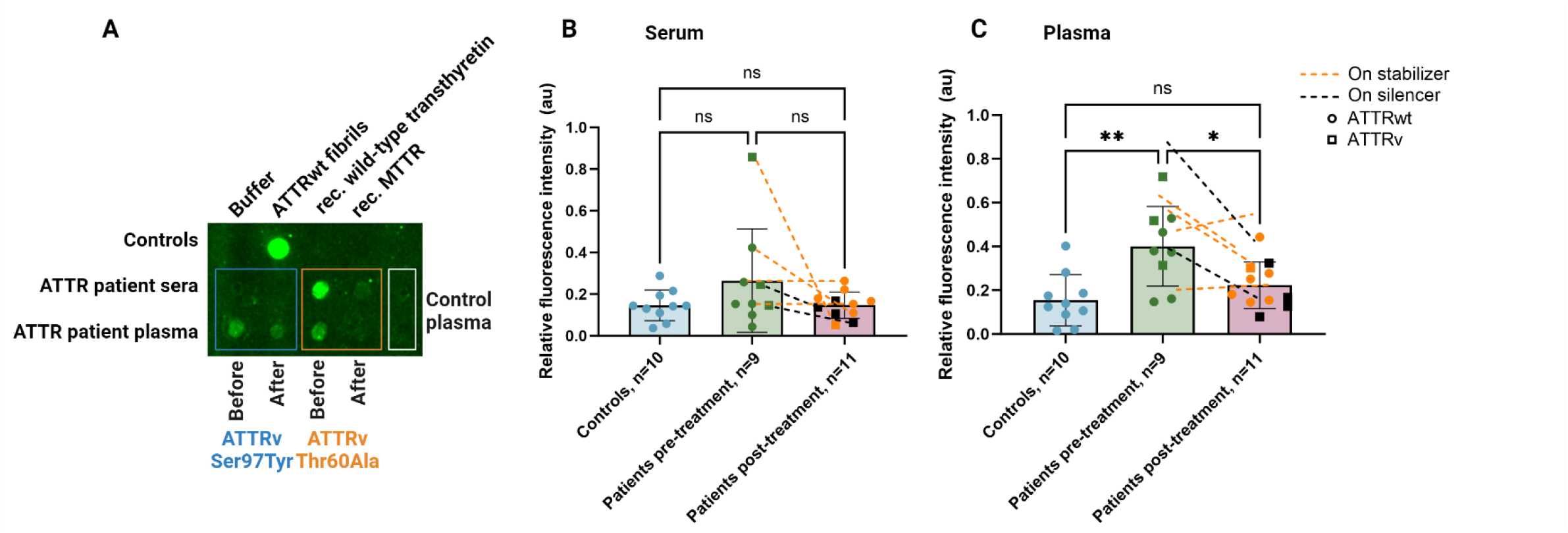
TAD1 detects unique ATTR species in plasma but not serum. **A**, Dot blotting of recombinant protein controls and extracted ATTRwt fibrils (top row), serum (middle row) and plasma (bottom row) from ATTRv amyloidosis patients before and after treatment with protein expression silencers, and control individuals. **B**, Quantification of TAD1 fluorescence intensity of binding to serum samples. **C**, Quantification of TAD1 fluorescence intensity of binding to plasma samples. Sample numbers are included in the labels. Each dot represents the average of three technical replicates from the same patient. Graphs include means (represented by bar heights) and standard deviation (represented by error bars). ns=not significant, * p<0.05, ** p <0.005.

Motivated by our discovery, we sought to characterize these ATTR species present in plasma. Since TAD1 binds ATTR fibrils and aggregates with high affinity^10^ we hypothesized whether these plasma species were also large in nature. We tested this hypothesis using plasma samples from ATTR amyloidosis patients in two complementary assays. First, we subjected ATTRwt amyloidosis, ATTRv amyloidosis, and control plasma samples to filtration using a 0.22 µM pore filter and used dot blotting to measure TAD1 binding to species in the filtrate and the void (Figure 4A). We found that TAD1 binds large ATTR species that cannot pass through the filter (Figure 4B).

**Figure 4.**
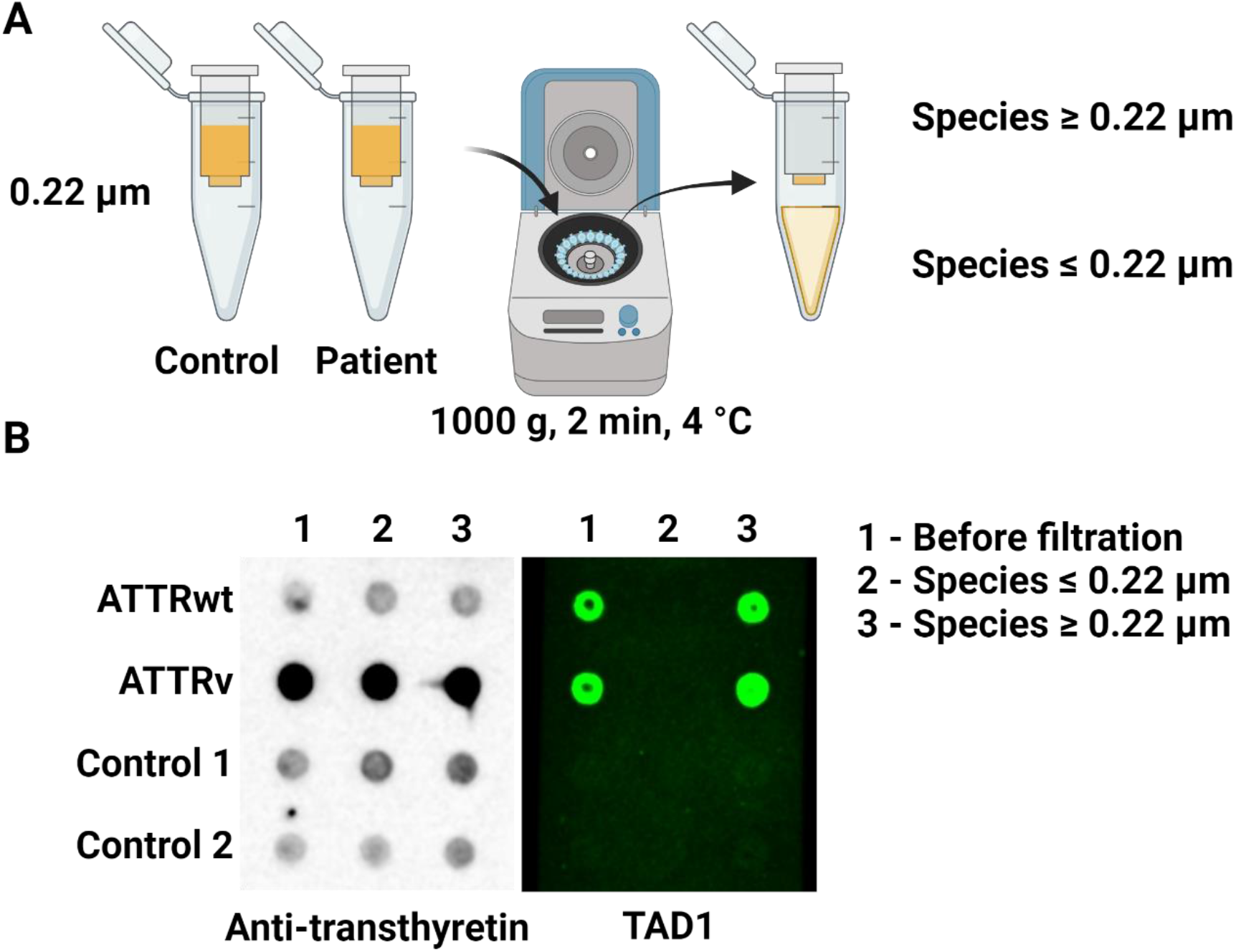
TAD1 binds large ATTR species in ATTR amyloidosis patient plasma. **A**, Scheme for filtration experiment. **B**, Filtered plasma of ATTR amyloidosis patients and controls probing with an anti-transthyretin antibody (left) and TAD1 (right).

In a complementary assay, we carried out a non-denaturing protein–protein band shift experiment, which allows us to visualize changes in electrophoretic behavior of a protein (soluble or aggregated) upon binding with a second protein (Figure 5). Briefly, we incubated ATTRwt amyloidosis patient plasma or control plasma with increasing concentrations of TAD1, ran the samples on a non-denaturing gel, and probed with an anti-transthyretin antibody. This experiment led to four key observations. The first observation from this experiment was the presence of oligomeric ATTR species in patient plasma, and the absence of these species in control plasma (Figure 5A-C). Second, upon binding to TAD1, these oligomers and tetrameric soluble transthyretin disappeared in a concentration-dependent manner. Third, we observed the concurrent accumulation of high molecular weight species that do not run through the gel (Figure 5A, B). Finally, we found that, in control plasma, there is an increase in tetrameric soluble transthyretin with increasing concentrations of TAD1 (Figure 5A, D). Together, these results indicate that there are large aggregated species present in the plasma of ATTR amyloidosis patients that can be detected by TAD1 in a conformation-dependent manner.

**Figure 5.**
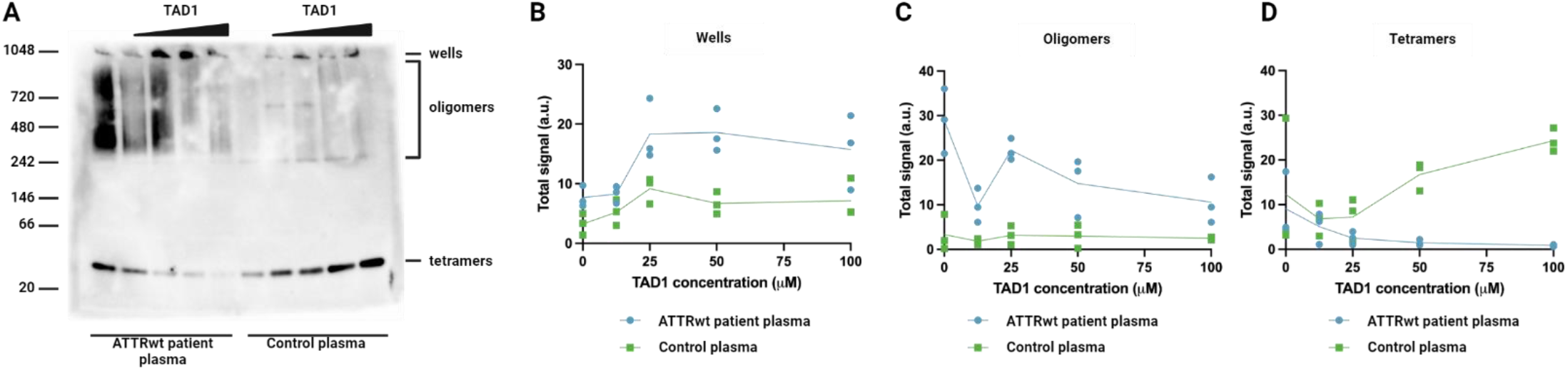
Gel shift assay of ATTRwt patient plasma and control plasma with increasing concentrations of TAD1, probed with an anti-transthyretin antibody. **A, B-D**, Quantification of transthyretin signal from gel shift assay in wells (left), oligomers (middle), and tetramers (right). All replicates are included.

## Discussion

In the present study, we use the cryogenic electron microscopy structures of ATTR fibrils to design a peptide (TAD1) that specifically binds large ATTR species (Figure 1). TAD1 targets aggregation-driving segments of transthyretin at the tips of fibrils as well as along the fibril surface in a conformation-dependent manner (Figures 1, 2, and Supplemental Figure 1). TAD1 detects not only ATTR fibrils after purification or in tissue homogenates, but also unique ATTR species in plasma of ATTR amyloidosis patients (Figures 3 and 4)^10^. We have determined these species to be high molecular weight oligomers that are distinct from ATTR oligomers found in neuropathic ATTR amyloidosis patients.

Other groups have identified non-native transthyretin (NNTTR) species in plasma of neuropathic patients using novel peptides and antibodies that target a specific transthyretin segment^3,4^. These species are different from the large ATTR species that we identified in plasma since the former are only present in neuropathic patients whereas the latter are found in our convenience sampling of both ATTRwt and ATTRv amyloidosis patients with cardiomyopathy or mixed phenotypes ^3,4,10^ Combined with previous results, our study implies that there may be multiple types of disease-associated transthyretin species in the blood of ATTR amyloidosis patients, including misfolded monomers and high molecular weight insoluble aggregates^3,4,10^ (Figures 3, 4, 5). Our study has uncovered a novel form of ATTR species present in cardiac ATTR amyloidosis patients.

An important query that results from this work is about the origin of ATTR aggregates observed in plasma. Oligomeric species can result from *de novo* nucleation of misfolded precursor proteins or from “shredding” of fibrillar deposits in tissues^13^. Previous observations suggest that ATTR aggregates in plasma may result from *de novo* nucleation. First, our TAD1 probe detects ATTR aggregates in asymptomatic carriers of pathogenic transthyretin alleles who have no detected pathology in the heart^10^. This finding suggests that the aggregation of transthyretin in the blood may identify amyloid pathophysiology before disease onset. On the other hand, there is a decrease of soluble transthyretin in patient serum that correlates with disease progression. This perhaps suggests that circulating transthyretin converts into insoluble species that are no longer detected in serum but can be detected in plasma^14^. If the plasma ATTR aggregates result from shredding of fibrillar deposits, the concentration of soluble transthyretin should not vary. However, we cannot rule out the possibility of both processes occurring concurrently after tissue deposition. This is especially important to consider because amyloid seeding is proposed to drive later stages of disease^6-8^. Perhaps there is a balance between both processes; that is, *de novo* nucleation may drive the formation of ATTR aggregates prior to deposition whereas the shredding of fibrils may contribute to the formation of ATTR aggregates in later stages of disease. Longitudinal studies of the aggregation of transthyretin in the blood of asymptomatic carriers of pathogenic alleles or other individuals at risk of developing ATTR amyloidosis would certainly begin to answer this question.

The biological significance of aggregates in plasma of ATTR amyloidosis patients has yet to be fully understood. The level of NNTTR species detected in neuropathic patients are reduced in patients under treatment with the tetramer stabilizer tafamidis, even when the cardiac deposition remains unchanged^3,4^. This observation suggests that oligomers may inflict physiological toxicity unrelated to cardiac amyloid deposition. Consistently, transthyretin oligomers obtained from recombinant sources were found toxic to cells *in vitro*^15,16^. Our experiments suggest that plasma ATTR aggregates are large oligomeric species, but it is unknown whether these aggregates are cytotoxic to cells. They could also be recognized and degraded by an individual’s immune system due to their altered conformation. A previous observation in a small cohort of cardiac ATTRwt amyloidosis patients with naturally occurring anti-amyloid antibodies and spontaneous reversal of ATTR-related cardiomyopathy alludes to the role of the immune system in ATTR amyloidosis prognosis^17^. The study of these plasma ATTR aggregates and their implications in immune response and cytotoxicity has the capacity to inform about the pathobiology of ATTR amyloidosis.

ATTR aggregates and their unique structures may constitute a novel target for clinical development. Other groups have developed reactive peptides and antibodies for specific detection of systemic amyloids that are structurally distinct from their native precursor proteins^18-20^. These tools have been successful in detecting amyloid deposits *in vivo*. Our TAD1 assay is a structure-guided approach that distinguishes between native and amyloidogenic transthyretin in blood. Early data suggest that TAD1 may be a promising biomarker for the detection and prediction of ATTR amyloidosis^10^.

In summary, we have used the structures of ATTR fibrils to design a novel peptide that binds *ex-vivo* ATTR fibrils, transthyretin aggregates generated *in vitro*, and ATTR species in plasma of ATTR amyloidosis patients. We have determined these species to be high molecular weight aggregates. Our findings open many questions about the biology and pathogenesis of ATTR amyloidosis and may lead to the development of novel therapeutic targets.

## Acknowledgements

The completion of this work would not be possible without many individuals. We are eternally grateful to the patients who generously donated their organs and tissues for our research. We thank the late Dr. Merrill Benson and the Indiana University, the Dallas Heart Study, and OHSU for providing patient samples. We thank the Electron Microscopy Core Facility at UT Southwestern, the Diamond lab for providing tau fibrils and brain lysate, and all members of the Saelices lab for their insightful discussion and feedback on experimental design and interpretation of results. Many thanks to the funding that makes this work possible.

