## Supplemental Information for "Structure-based probe reveals the presence of large transthyretin aggregates in plasma of ATTR amyloidosis patients"

**Supplemental Table 1. Third generation peptide inhibitor sequences.**

| Peptide name | Sequence |
|---|---|
| TAB3-01 | RRRRHVAHPFVEFTE |
| TAB3-02 | RRRRSYVTNPTSYAVT |
| TAB3-03 | RRRRHVAHPFVEFTERRRRSYVTNPTSYAVT |
| TAB3-04 | RRRRSYVTNPTSYAVTRRRRHVAHPFVEFTE |
| TAB3-05 | RRRRHVAHPFV(n-me)EFTERRRRSYVTNPTSY(n-me)AVT |
| TAB3-06 | RRRRSYVTNPTSY(n-me)AVTRRRRHVAHPFV(n-me)EFTE |
| TAB3-07 | RRRRHVAHPFVEFTEGSRRRRSYVTNPTSYAVT |
| TAB3-08 | RRRRHVAHPFVEFTEGGSRRRRSYVTNPTSYAVT |
| TAB3-09 | RRRRHVAHPFVEFTEGGGSRRRRSYVTNPTSYAVT |
| TAB3-10 | RRRRHVAHPFVEFTEGGGSTRRRRSYVTNPTSYAVT |
| TAB3-11 | RRRRHVAHPFVEFTEGGGSARRRRSYVTNPTSYAVT |
| TAB3-12 | RRRRHVAHPFVEFTEGGGSTERRRRSYVTNPTSYAVT |
| TAB3-13 | RRRRHVAHPFVEFTEGGGSAERRRRSYVTNPTSYAVT |

**Supplemental Table 2. Sequences of peptide detection probes.**

| Peptide name | Sequence | N-Terminal Modification | N-Terminal Epitope Tag |
|---|---|---|---|
| TAD1 | HHHHHHRRRRHVAHPFVEFTEGGGSTERRRRSYVTNPTSYAVT | FITC-Aminohexanoic acid (Ahx) | Polyhistidine |
| TAD2 | YPYDVPDYARRRRHVAHPFVEFTEGGGSTERRRRSYVTNPTSYAVT | FITC-Ahx | Hemagglutinin |
| TAD3 | DYKDDDDKRRRRHVAHPFVEFTEGGGSTERRRRSYVTNPTSYAVT | FITC-Ahx | FLAG |

**Supplemental Table 3. List of ATTR amyloidosis cardiac tissue samples included in the study.**

| Genotype | Origin | Sex | Age at collection | Neuropathy signs |
|---|---|---|---|---|
| ATTRwt | Autopsy | Male | 84 | No |
| ATTRv-V30M | N/A | N/A | N/A | N/A |
| ATTRv-V122I | Autopsy | Male | N/A | N/A |

**Supplemental Table 4. Characteristics of ATTR amyloidosis plasma samples included in the study.**

| Clinical Characteristics | Controls (n=10) | Pre-Treatment Patients (n=9) | Post-Treatment Patients (n=11) |
|---|---|---|---|
| **Type of ATTR amyloidosis** | | | |
| Wild-type, n (%) | N/A | 6 (66.7) | 6 (54.5) |
| Variant, n (%) | N/A | 3 (33.3) | 5 (45.5) |
| **Variant Types** | | | |
| ATTR-V30M (*p.V50M) | N/A | 1 (11.1) | 1 (9.1) |
| ATTR-T60A (*p.T80A) | N/A | 1 (11.1) | 3 (27.3) |
| ATTR-S97Y (*p.S117Y) | N/A | 1 (11.1) | 1 (9.1) |
| **Treatment Type** | | | |
| Transthyretin tetramer stabilizer | N/A | N/A | 7 (63.6) |
| Transthyretin mRNA silencer | N/A | N/A | 4 (36.4) |

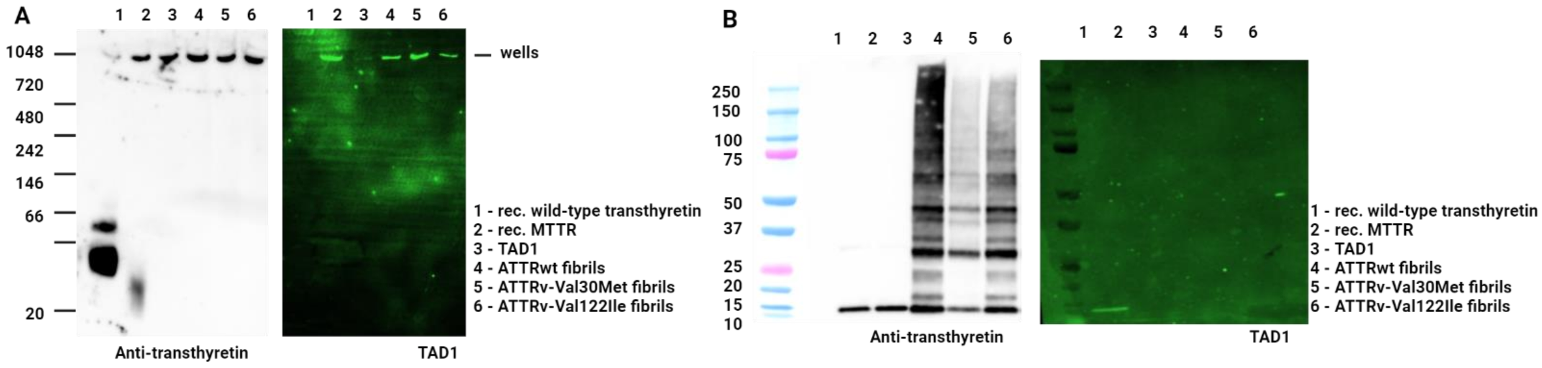


**Supplemental Figure 1. TAD1 binds ATTR fibrils in a conformation-dependent manner. A**, Native gel electrophoresis of TAD1, recombinant transthyretin and amyloid extracted from ATTR amyloidosis patients, blotted and probed with an anti-transthyretin antibody (left) and TAD1 (right). **B**, Gel electrophoresis of same samples under denaturing conditions, blotted and probed with an anti-transthyretin antibody (left) and TAD1 (right).

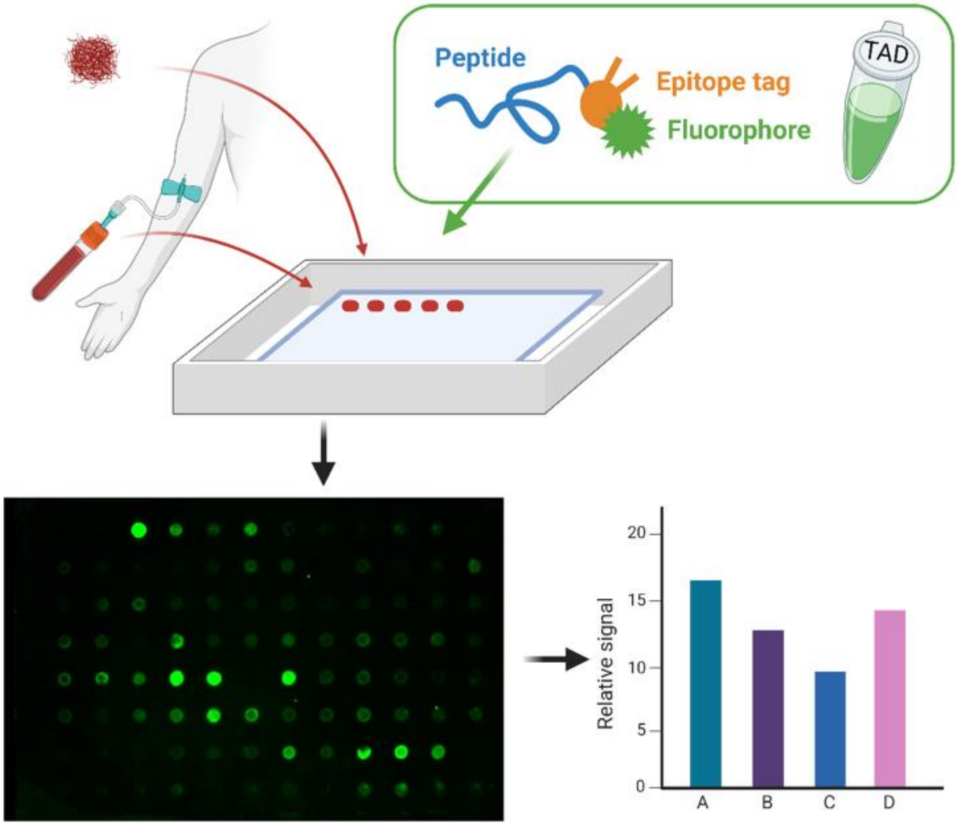


**Supplemental Figure 2. Dot blotting allows for quantification of TAD binding to samples of interest.** Recombinant protein or patient samples (extracted fibrils from heart, cardiac lysates, or blood) are applied to nitrocellulose membrane then incubated with TAD peptide. TAD binding to sample is measured through relative fluorescence intensity, which can then be quantified to provide insight into the relative fibrillar content of each sample.

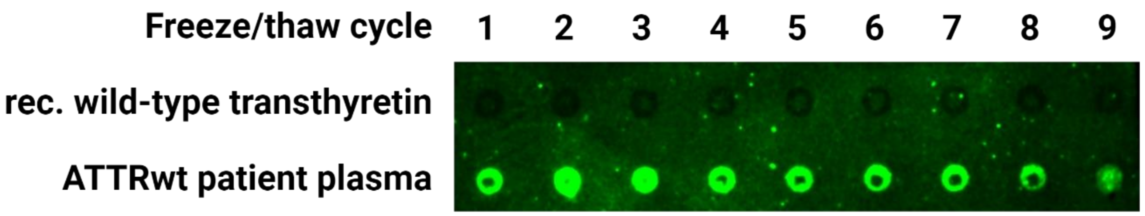


**Supplemental Figure 3. TAD1 binding is not impacted by freeze/thaw cycles.** Dot blotting of recombinant wild-type transthyretin and ATTRwt patient plasma after eight freezing and thawing cycles shows no remarkable difference in TAD1 binding to plasma samples after at least 6 cycles.
